# InClust+: the multimodal version of inClust for multimodal data integration, imputation, and cross modal generation

**DOI:** 10.1101/2023.03.13.532376

**Authors:** Lifei Wang, Rui Nie, Yankai Cai, Anqi Wang, Hanwen Zhang, Jiang Zhang, Jun Cai

## Abstract

With the development of single-cell technology, many cell traits (e.g. gene expression, chromatin accessibility, DNA methylation) can be measured. Furthermore, the multi-omic profiling technology could jointly measure two or more traits in a single cell simultaneously. In order to process the various data accumulated rapidly, computational methods for multimodal data integration are needed. Previously, we developed inClust, a flexible all-in deep generative framework for transcriptome data. Here, we extend the applicability of inClust into the realm of multimodal data by adding two mask modules: an input-mask module in front of the encoder and an output-mask module behind the decoder. We call this augmented model inClust+, and apply it to various multimodal data. InClust+ was first used to integrate scRNA and MERFISH data from similar cell populations and to impute MERFISH data based on scRNA data. Then, inClust+ is shown to have the capability to integrate a multimodal data contain scRNA and scATAC or two multimodal CITE datasets with batch effect. Finally, inClust+ is used to integrate a monomodal scRNA dataset and two multimodal CITE datasets, and generate the missing modality of surface protein in monomodal scRNA data. In the above examples, the performance of inClust+ is better than or comparable to the most recent tools to the corresponding task, which prove inClust+ is a suitable framework for handling multimodal data. Meanwhile, the successful implementation of mask in inClust+ means that it can be applied to other deep learning methods with similar encoder-decoder architecture to broaden the application scope of these models.

## Introduction

Recently, the progress of single-cell technology makes it possible to obtain a variety of traits in a single cell. Single-cell RNA sequencing (scRNA-seq), the pioneer of single cell technology, measures the gene expression level of a single cell[1, 2]; single-cell assay for transposase-accessible chromatin using sequencing (scATAC) profile chromatin accessibility in a single cells[3]; single-cell bisulfite sequencing (scBS-seq) could reveal genome-wide DNA methylation status at the single cell level[4]. Various single-cell methods have greatly promoted our understanding of cells. As a result, the heterogeneity of cell population was revealed[2, 5], the trajectory of cell development was inferred[6], and the gene regulatory network for controlling expression program was reconstructed[7]. But data collected in one modality of single cell just represent a limited side view of the cell state. In order to obtain more holistic and comprehensive information, data from different modalities need to be integrated together to better reveal the biological significance of the data.

Initially, the integration of data from different modalities of single cell was accomplished by computational approach[8, 9]. Then, Multi-omic profiling technology that could jointly profile multiple traits in a single cell is developed[10]. Several methods (e.g. SNARE-seq[11], sci-CAR[12], Paired-seq[13], and SHAER-seq[14]) could simultaneously measure the gene expression and chromatin accessibility in a single cell. Cellular indexing of Transcriptomes and Epitopes by sequencing (CITE-seq) could jointly profile the gene expression and a panel of cell surface protein[15, 16]. The scNMT-seq could profile chromatin accessibility, DNA methylation, and transcription in single cells at one time[17].

Several computational approaches have been developed to process and integrate data in single-cell analysis. Some are universal methods that could handle situations with either multiple monomodal data profiled from different cells in similar populations or multimodal data extracted from single cells[18]. Others are specially designed for processing data generated by multi-omics profiling technology[19, 20]. Usually, for data generated by multi-omics profiling technology, the information from different modalities is coded by different encoders first, and then integrated in the latent space. The one-to-one correspondence between encoders and modalities is mainly due to the fact that data from different modalities have different data formats and lengths, which can’t be encoded by the same encoder. Following the encoding, the coded information from different modalities is integrated in the latent space, through adversarial loss[21], mixture of expert[22], attention-transfer[23], regulatory interaction information[24], and so on. After integrating data from different modalities, data from one modality could be used to impute data from another modality, which may have a low resolution that needs to be enhanced[18]. Meanwhile, the integration of multimodal data makes the translation between different modalities possible[21]. Furthermore, the multimodal data could be used as a reference to generate data on the missing modality in monomodal data[25].

Previously, we presented inClust (integrated clustering), a flexible all-in deep generative framework with functional modules for embedding auxiliary information, latent space vector arithmetic, and clustering[26]. It could complete the task of conditional out-of-distribution generation in supervised mode, transfer labels and filter out new cell type in semi-supervised mode, and identify spatial domains in unsupervised mode. Here, we extend the inClust by adding two new modules, namely, the input-mask module in front of encoder and the output-mask module behind decoder. We named our augmented inClust as inClust+, and demonstrated that it could integrate multimodal data profiled from different cells in similar populations or from a single cell. Furthermore, the capability of inClust+ to cross modal imputation and generation is also demonstrated. All results show that inClust+ is an ideal tool to deal with multimodal data, and adding masks is a suitable way to model augmentation in the field of multi-omics.

## Results

### InClust+ imputes genes for MERFISH data after integrating scRNA and MERFISH data

The rationale for integrating scRNA and MERFISH data by inClust+ is simple: just treat the multimodal data from different modalities as scRNA data from different batches. In addition, the input-mask module and the output-mask module would enable the gene imputation in MERFISH data based on transferring knowledge from scRNA data (Fig 1A). For scRNA data, inClust+ uses the common genes to reconstruct common genes and scRNA-specific genes for scRNA data. The reconstructed expression profile of the common genes and the scRNA-specific genes were compared with the real expression profile to update the model parameters (Fig 1B). For MERFISH data, only reconstructed expression profile of common genes is used for parameter updating (Fig 1C). Although expression of scRNA-specific genes in MERFISH data did not contribute to the updating of model parameters, they were still reconstructed as a by-product. Since the encoder and decoder used for scRNA and MERFISH data are the same, the reconstructed expression of scRNA-specific genes in MERFISH data could depend on the knowledge transferred from scRNA data (Fig 1A).

**Figure 1.**
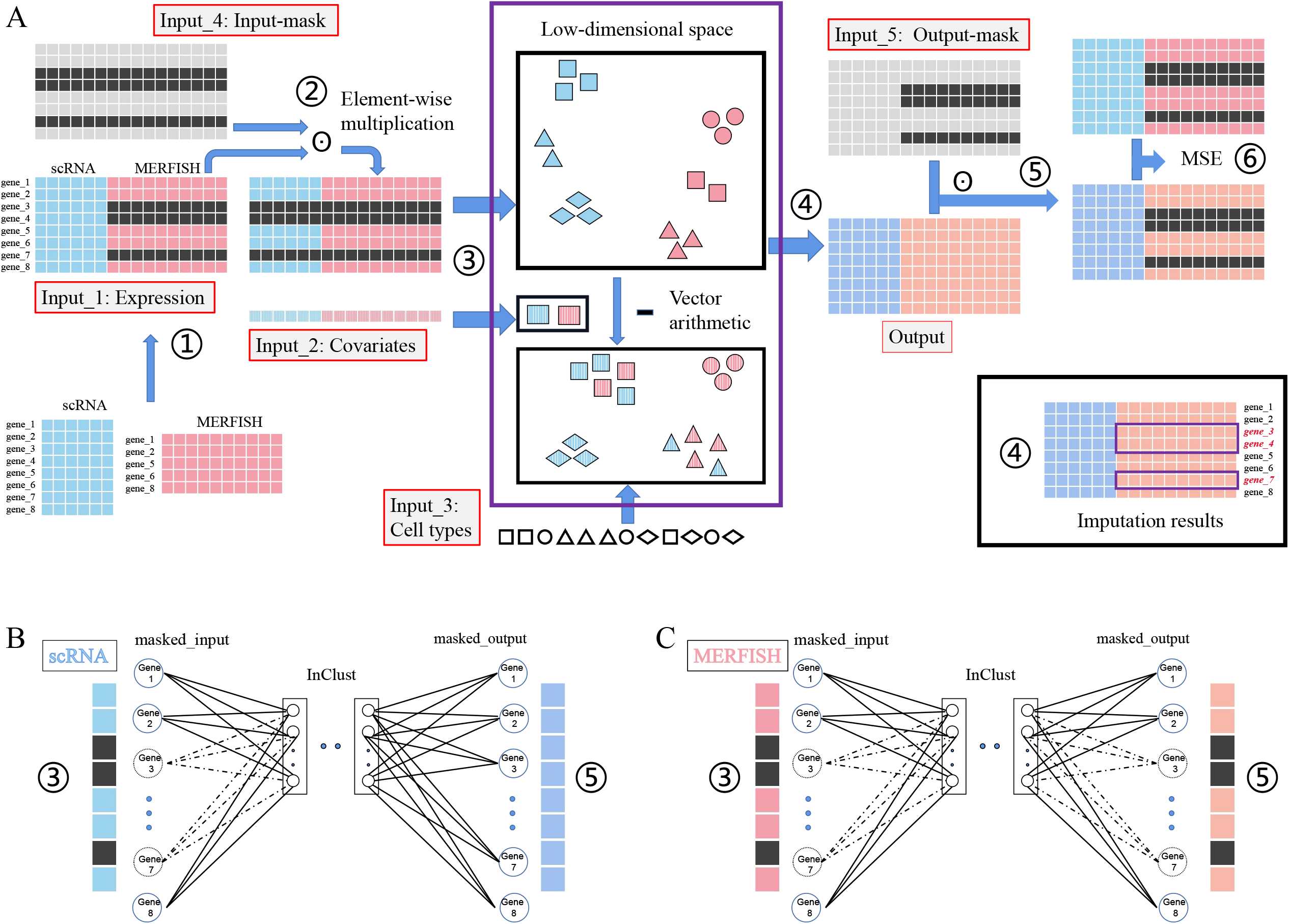
The diagram for integration of multiple monomodal (unpaired) data and subsequently gene imputation by inClust+. A. The workflow of inClust+ for integration of scRNA and MERFISH data, and subsequently gene imputation for MERFISH data. Training: ①Generation of the training dataset. To form a training dataset, the data from scRNA and MERFISH were aligned with common genes, and the missing scRNA-specific genes in MERFISH data were filled with 0. ②Generation of the masked-input for the encoder in inClust+. The training dataset multiplies element-wise with an input-mask matrix. The input-mask matrix is as large as the training dataset. In the input-mask matrix, the positions of common genes are filled with 1, and the positions of scRNA-specific genes are filled with 0. The result of multiplication is the masked-input of the encoder, and only common gene is effective. ③Data encoding, covariates elimination and data integration. Firstly, the expression profiles of common genes are encoded in low-dimensional space, and then the covariates (modalities) are eliminated by subtracting covariates information from expression profile, and the data from different modalities are integrated through the constraints of the cell type information. ④ Reconstruction of expression profile for both common genes and scRNA-specific A. genes. The decoder outputs the expression profile of the common and scRNA-specific genes. ⑤Generation of the masked-output for loss calculation. The output multiplies element-wise with an output-mask matrix, which is as large as the output. In the output-mask matrix, the place of scRNA-specific genes in MERFISH filled with 0 and the other place filled with 1. ⑥Calculation of the loss for backpropagation. The mean squared error (MSE) between masked-outputs and the training data is calculated as the loss. Imputation: After training, the output of the decoder (step ④) would impute the missing scRNA-specific genes in MERFISH data. B. Training inClust+ with scRNA data. In encoder, only the expression data of common genes are the effective inputs. So, in the first layer of the encoder, only the corresponding connections actually contribute to the encoding process. In decoder, both common genes and scRNA-specific genes are reconstructed and pass through the mask. Then the loss between input and output with both the common genes and specific genes is calculated, and all connections in the last layer contribute to the loss. In short, when training with scRNA data, inClust+ uses common genes to reconstruct common genes and scRNA-specific genes. C. Training inClust+ with MERFISH data. In encoder, only the expression data of common genes are the effective inputs. So, in the first layer of the encoder, only the corresponding connections actually contribute to the encoding process. In decoder, both common gene and scRNA-specific gene are reconstructed, while the scRNA-specific genes are filtered out by the output-mask. Loss is calculated according to the common genes, so only connections corresponding to common genes in the last layer of decoder contribute to the calculation of loss. In short, when training with MERFISH data, inClust+ use common genes to reconstruct common genes. However, after training, inClust+ would output common genes and scRNA-specific genes from the input of common genes.

For comparison, we randomly selected 80% of genes in MERFISH data as common genes, and the rest as test genes (scRNA-specific) waiting for imputations, as described in the uniPort[18]. The inClust+ first encodes the scRNA and MERFISH data into latent space respectively. As the input data (Fig 2A), the encoded representations from different modalities are also separated in the latent space (Fig 2B). After covariates (modalities) removal by vector subtraction, the samples from different modalities were mixed together and clustered according to their cell types (Fig2 C). As the uniPort[18], the evaluation of imputation is calculated using median and average Spearman correlation coefficients (mSCC and aSCC), and the median and average Pearson correlation coefficients (mPCC and aPCC) over imputed and true testing genes. As shown in the plot, inClust+ demonstrated the higher mSCC (0.243), aSCC (0.255), mPCC (0.263), and aPCC (0.322), above those of uniPort (mSCC of 0.236, aSCC of 0.247, mPCC of 0.233, and aPCC of 0.274) (Fig. 2D).

**Figure 2.**
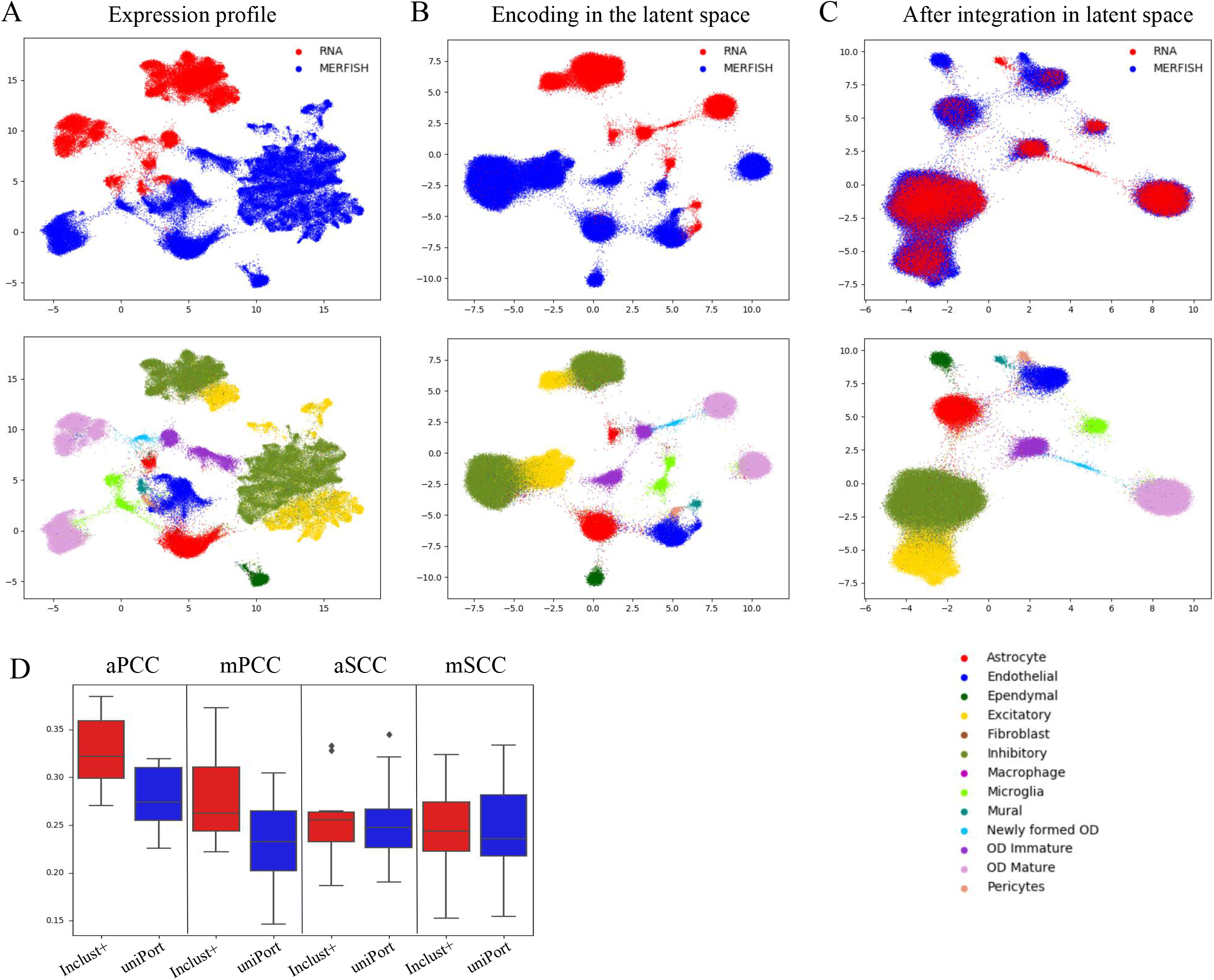
The results for integration of multiple monomodal (unpaired) data and subsequently gene imputation by inClust+. A. The UMAP plot of the scRNA and MERFISH data (the top 50 PCs) colored by the modalities (top) and cell types (bottom). B. The UMAP plot of the low dimensional representations with covariate effects for the scRNA and MERFISH data in inClust+ colored by the modalities (top) and cell types (bottom). C. The UMAP plot of the low dimensional representations without the covariate effects for the scRNA and MERFISH data in inClust+ colored by the modalities (top) and cell types (bottom). D. Comparison of imputation capability of inClust+ and uniPort. Boxplots of average and median Pearson correlation coefficients (aPCC and mPCC) (n = 12), and average and median Spearman correlation coefficients (aSCC and mSCC) (n = 12) between real and imputed MERFISH genes generated by inClust+ and uniPort were plotted.

### InClust+ integrate multi-omics dataset

Single cell multi-omics can extract information from different cellular components of a single cell at the same time, with different data types and lengths. Those multimodal data contain more information than monomodal data, and need more specific methods to process, rather than integrating multiple monomodal data. By flexibly adjusting the input (output) mask modules, inClust+ can be transformed into a model specially used for multimodal data processing. In order to use data from both modalities at the same time, data from both modalities are stacked together in the input (Fig 3A). Accordingly in the model, the first layer of encoder could be regarded as two independent neural network layers stacked together, and each part corresponds to data from one modality (Fig 3B) (e.g. one for scRNA and the other for surface protein or scATAC). The last layer of the decoder is also divided into two parts, respectively, which reconstruct the data of different modalities. Each round of the model training is divided into four stages. In the first two phases (Fig 3B, C), inClust+ uses data from one modality to reconstruct data from corresponding modality (e.g. in the first stage, inClust+ uses the surface protein data to rebuild surface protein data. In the second phase, inClust+ uses the scRNA data to reconstruct scRNA data). On the contrary, in the last two phases (Fig 3D, E), inClust+ uses data from one modality to reconstruct data from another modality (e.g. in the third phase, inClust+ uses the surface protein data to reconstruct scRNA data. In the fourth phase, inClust+ uses the scRNA data to reconstruct surface protein data). The rationale for four stages training is as follows: in the first and second stages, the upper and lower part of the first layer of the encoder are coupled with the corresponding part of the last layer of the decoder. This could be thought of as an encoder/decoder combination for data from one modality. Each encoder/decoder combination updated themselves relatively independently. In contrast, in the third and fourth stages, the upper and lower part of the first layer of the encoder are coupled with the opposite part of the last layer of the decoder. This is an attempt to translate between different modalities in a single cell and integrate them more toughly. Furthermore, the batch effect between different datasets could be explicitly removed by vector arithmetic in the latent space as the original inClust (Fig 3A).

**Figure 3.**
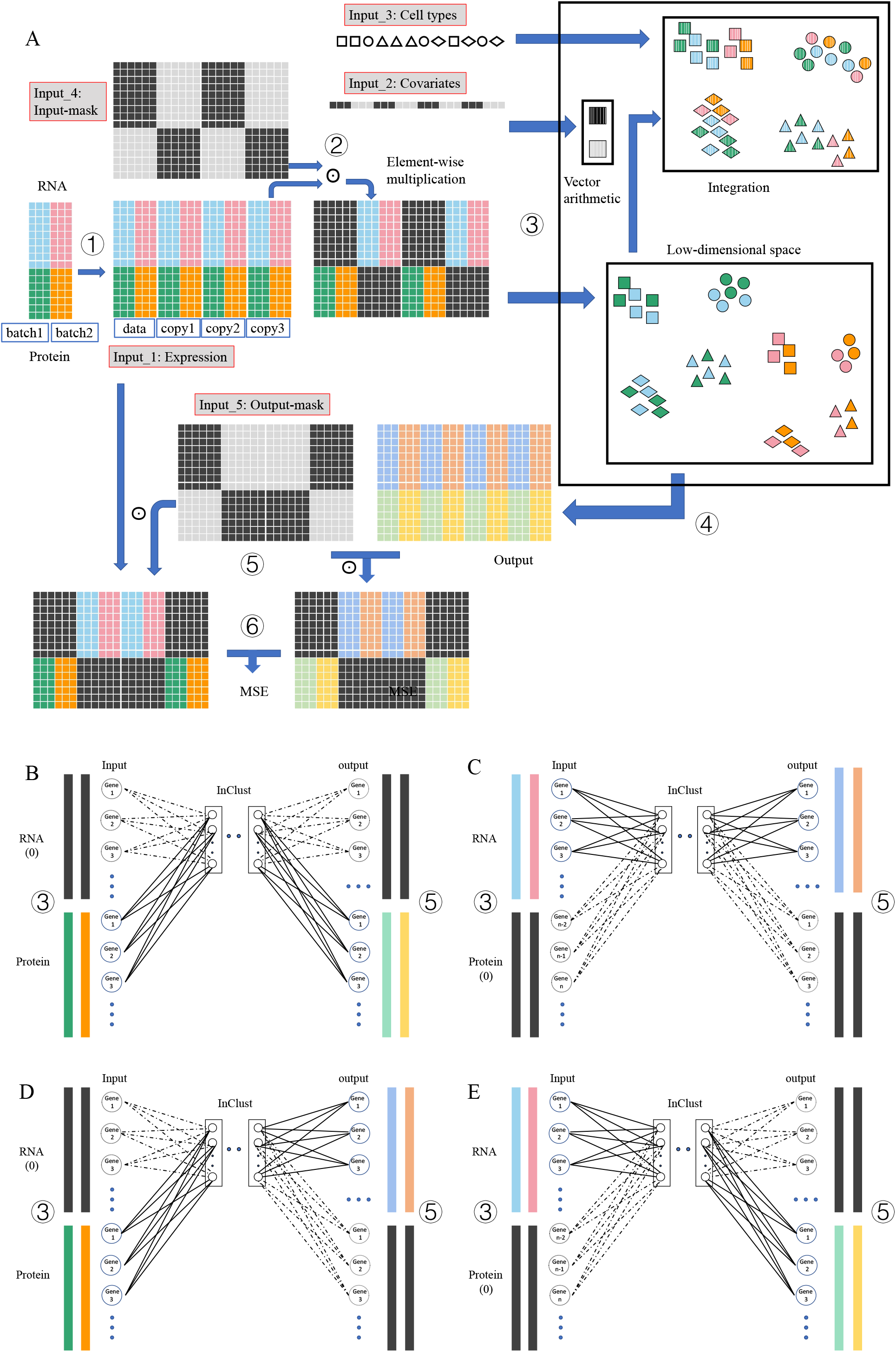
The diagram for integration of multimodal (paired) datasets by inClust+ A. The workflow of inClust+ for integration of multimodal (paired) datasets (scRNA and protein). Training: ①Generation of the training dataset. To form a training dataset, the multimodal (paired) datasets were duplicated 3 times, and concatenated together. ②Generation of the masked-input for the encoder in inClust+. The training dataset multiplies element-wise with an input-mask matrix. The input-mask matrix is as big as the training dataset, and could be equally divided into four parts, each part being as big as the original multimodal (paired) datasets. In the first and the third part, the positions of scRNA data are filled with 0, and the positions of protein data are filled with 1. Alternatively, in the second and the fourth part, the positions of scRNA data are filled with 1, and the positions of protein data are filled with 0. The result of multiplication is the masked-input for the encoder, with alternate training data of scRNA and protein. ③Data encoding, covariates elimination and data integration. The data from different modalities (scRNA or protein) in the masked-input are encoded by the different parts of encoder into the low-dimensional space, and integrated through the constraints of the cell type information. The batch effect is removed by vector subtraction in latent space. ④Reconstruction for both scRNA data and protein data. The decoder simultaneously outputs the reconstructed scRNA data and the reconstructed protein data. ⑤Generation of the masked-output for loss calculation. The output multiplies element-wise with an output-mask matrix. The output-mask matrix is as large as the output, and could be equally divided into four parts, each part being as big as the original multimodal (paired) datasets. In the first and the fourth part, the positions of scRNA data are filled with 0, and the positions of protein data are filled with 1. Alternatively, in the second and the third part, the positions of scRNA data are filled with 1, and the positions of protein data are filled with 0. The result of multiplication is the masked-output. ⑥Calculation of the loss for backpropagation. The MSE between masked-output and the masked-input is calculated as the loss. Data integration: After training, encoded low-dimensional representations are mixed together and clustered according to the cell types without the effect of covariate (batches and modalities). B. The first training phase. In this stage, due to the influence of mask, only protein data is effective for input and output. Therefore, only the corresponding connections in the first layer (lower part) of the encoder and the last layer (lower part) of the decoder actually contribute to the training process. In short, inClust+ uses protein data to reconstruct protein data. C. The second training phase. In this stage, due to the influence of mask, only scRNA data is effective for input and output. Therefore, only the corresponding connections in the first layer (upper part) of the encoder and the last layer (upper part) of the decoder actually contribute to the training process. In short, inClust+ uses scRNA data to reconstruct scRNA data. D. The third training phase. In this stage, due to the influence of mask, only protein data is effective for input and scRNA data is effective for output. Therefore, only the corresponding connections in the first layer (lower part) of the encoder and the last layer (upper part) of the decoder actually contribute to the training process. In short, inClust+ uses protein data to reconstruct scRNA data. E. The fourth training phase. In this stage, due to the influence of mask, only scRNA data is effective for input and protein data is effective for output. Therefore, only the corresponding connections in the first layer (upper part) of the encoder and the last layer (lower part) of the decoder actually contribute to the training process. In short, inClust+ uses scRNA data to reconstruct protein data.

We first applied inClust+ to integrate the multimodal PBMC data with scATAC data and scRNA data (Fig S1). Before integration, scATAC data and scRNA data were separated in the original space (Fig S2A). After integration by inClust+, the data from scATAC and scRNA are mixed together in the latent space (Fig S2B). As in uniPort, the Batch Entropy score is used to measure the degree of mixing cells across datasets and the Silhouette coefficient is used to evaluate the separation of biological distinctions[18]. The result shows that inClust+ has obtained a Batch Entropy score of 0.686 and a Silhouette coefficient of 0.808, which is much higher than those of uniPort (Batch Entropy score of 0.619 and Silhouette coefficient of 0.64) (Fig S2C).

We then applied our model to integrate multiple multimodal datasets with some batch effect. Two CITE datasets from different donors with some batch effects are used. Both scRNA data (Fig4 A) and protein data (Fig4 B) in CITE datasets have batch effects. The inClust+ integrates data from different modalities in the latent space (Fig 4C). And the vector arithmetic further integrates data from different batches (Fig 4D).

**Figure 4.**
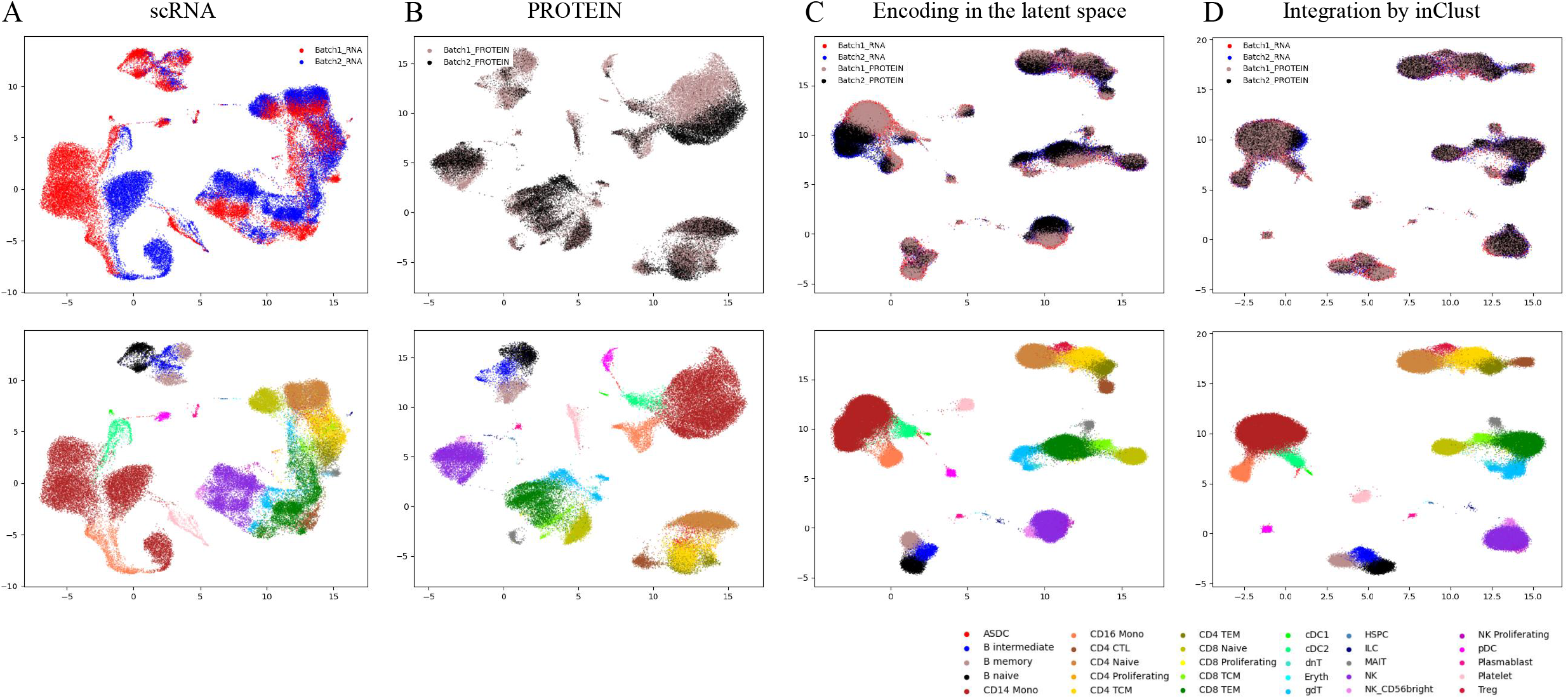
The results for integration of multiple multimodal (paired) datasets by inClust+. A. The UMAP plot of the scRNA data colored by the batches (top) and cell types (bottom). B. The UMAP plot of the Protein data colored by the batches (top) and cell types (bottom). C. The UMAP plot of the low dimensional representations with batch effects for the scRNA and Protein data in inClust+ colored by the covariate (top) and cell types (bottom). D. The UMAP plot of the low dimensional representations without the batch effects for the scRNA-seq and Protein data in inClust+ colored by the covariate (top) and cell types (bottom).

### Cross modal generation by InClust+

The multi-omics dataset contains data from multiple modalities, and could be used as a reference to complete the monomodal data into multimodal data. Our inClust+ can extract information from multi-omics reference, and translate monomodal data into data of another modality. As the situation for multimodal integration, the first layer of encoder and the last layer of decoder could be regarded as two independent neural network layers stacked together to handle the stack data of two modalities (Fig 5A). The translation from scRNA data into data of surface protein in the multimodal reference was carried out in two stages in each round of training. In the first stage, inClust+ uses the scRNA data to reconstruct scRNA data (Fig 5B). Alternatively, in the second phase, inClust+ uses the scRNA data to reconstruct protein data (Fig 5C). There is the third stage for the monomodal data that needs to be completed. In this stage, inClust+ uses the scRNA data to reconstruct scRNA data in the monomodal dataset (Fig 5B). After training, inClust+ could transfer labels from the scRNA data in the multimodal reference to the scRNA monomodal dataset. Meanwhile, based on the scRNA data in the monomodal dataset, the corresponding protein data could be generated by automatic translation.

**Figure 5.**
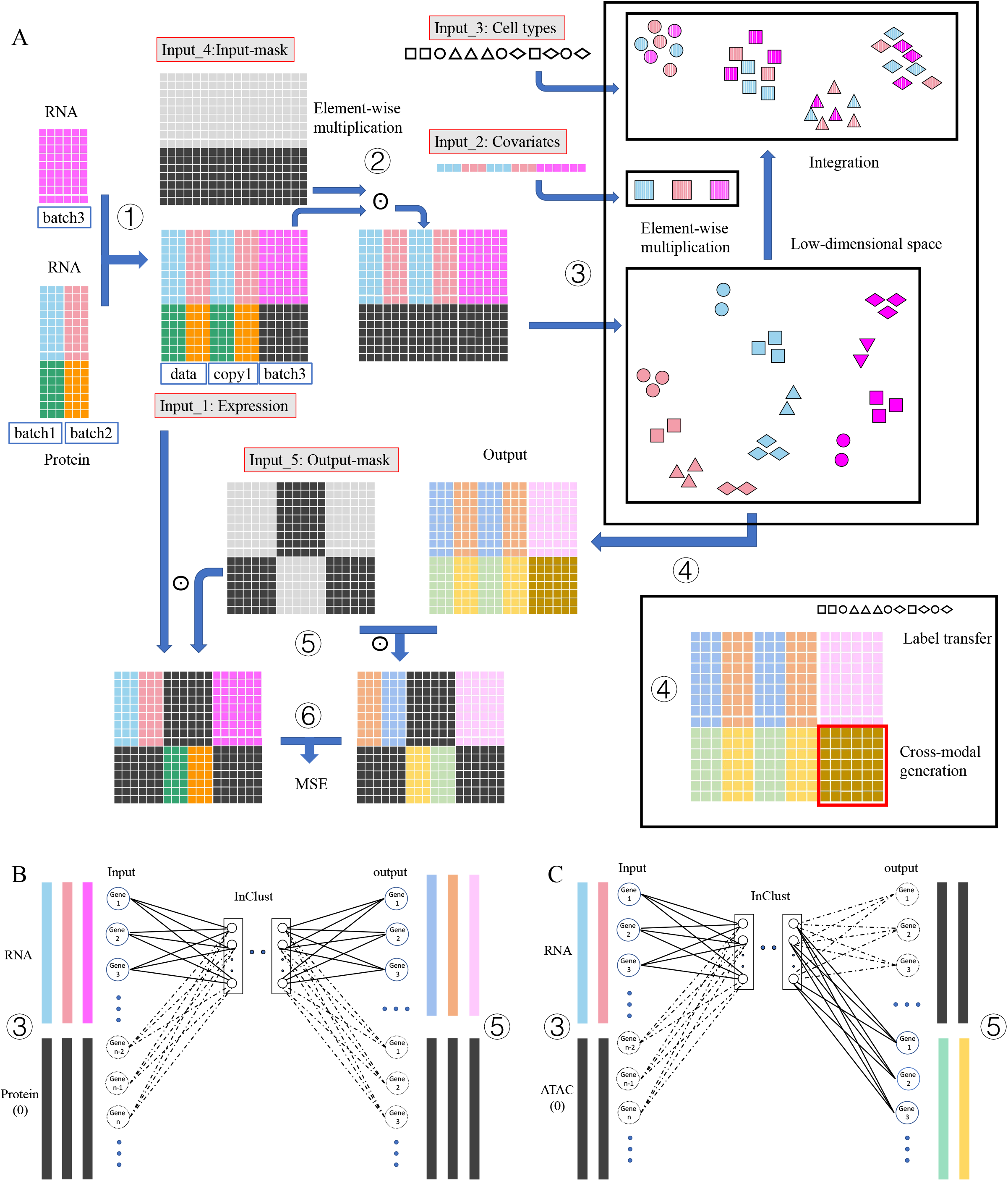
The diagram for cross modal generation of inClust+ A. The workflow of inClust+ for integration of two multimodal datasets and a monomodal scRNA dataset, and subsequently cross modal generation. Training: ① Generation of the training dataset. To form a training dataset, the data from scRNA and CITE were aligned with shared genes, and the missing Protein data in scRNA data were filled with 0. The CITE data were duplicated one time, and concatenated together. ②Generation of the masked-input for the encoder in inClust+. The training dataset multiplies element-wise with an input-mask matrix. The input-mask matrix is as large as the training dataset, and the positions of scRNA data are filled with 1, and the positions of Protein data are filled with 0. The result of multiplication is the masked-input of the encoder, and only scRNA data is effective. ③Data encoding, covariates elimination and data integration. Firstly, the expression profiles of scRNA are encoded in low-dimensional space, and then the batch (covariates) effect is eliminated by subtracting batch information from expression profile, and the data from different batches are integrated through the constraints of the cell type information and inherent biological characteristic. ④The decoder simultaneously outputs the reconstructed scRNA data and the reconstructed Protein data. ⑤ Generation of the masked-output for loss calculation. The output multiplies element-wise with an output-mask matrix. The output-mask matrix is as large as the output. In the output-mask matrix, the position corresponding to duplicated CITE data could be equally divided into two parts, each part being as big as one CITE data. In the first part, the positions of scRNA data are filled with 1, and the positions of Protein data are filled with 0. Alternatively, in the second part, the positions of scRNA data are filled with 0, and the positions of Protein data are filled with 1. Meanwhile, the positions of scRNA data in monomodal data are filled with 1, and the positions of Protein in monomodal data are filled with 0. The result of multiplication is the masked-output. ⑥Calculation of the loss for backpropagation. The MSE between masked outputs and the masked training dataset is calculated as the loss. Label transfer and cross modal generation: after training, the labels are transferred from cells of multimodal data to the cells of monomodal data in the same clusters. The output of the decoder (step ④) would generate the missing modality in monomodal data. B. Training inClust+ with scRNA data. In these stages, only scRNA data is effective for input and output. Therefore, only the corresponding connections in the first layer (upper part) of the encoder and the last layer (upper part) of the decoder actually contribute to the training process. In short, inClust+ uses scRNA data to reconstruct scRNA data. C. Training inClust+ with scRNA data and translating them into Protein data. In these stages, due to the influence of mask, only scRNA data is effective for input and Protein data is effective for output. Therefore, only the corresponding connections in the first layer (upper part) of the encoder and the last layer (lower part) of the decoder actually contribute to the training process. In short, inClust+ uses scRNA data to reconstruct Protein data.

We evaluated the capability of inClust+ to complete a monomodal dataset into a multimodal dataset through two CITE-seq references and a scRNA-seq test set. The UMAP plots show that inClust+ could integrate the scRNA data from different datasets well (Fig 6A, FigS3). And the results of the labels transferring are plotted in the confusion matrix, which show that inClust+ is better than sciPENN, with accuracy of 0.947 in inClust+ (Fig 6B) and 0.915 in sciPENN (Fig 6C). The generated protein expression profile by inClust+ was visualized by UMAP (Fig 6D). And the prediction accuracy of protein expression is measured by calculating the Pearson correlation and the Spearman correlation between the predicted data and real data. The results show that inClust+ (mSCC of 0.334, mPCC of 0.376) is comparable to sciPENN (mSCC of 0.356, mPCC of 0.405), which is specially optimized for protein prediction related to CITE multimodal data[25] (Fig 6E).

**Figure 6.**
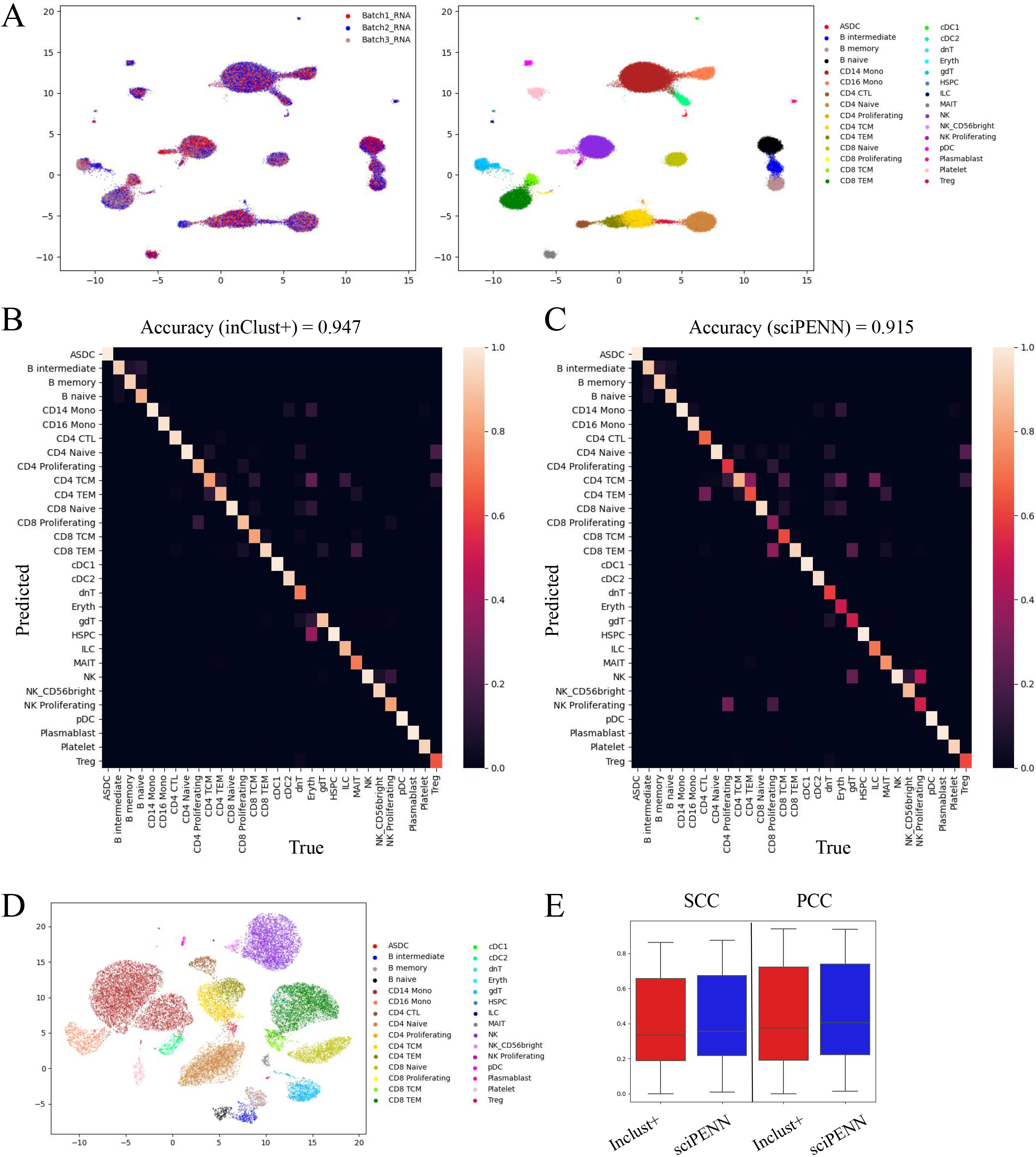
The results for cross modal generation by inClust+ A. The UMAP plot of the low dimensional representations without the covariate effects for the scRNA in inClust+ colored by the covariate (left) and cell types (right). B. Heatmap for the confusion matrix of results generated by inClust+ with average accuracy above. C. Heatmap for the confusion matrix of results generated by sciPENN with average accuracy above. D. The UMAP plot visualization of generated Protein data by inClust+ E. Comparison of cross modal generation results of inClust+ and sciPENN. Boxplots of Pearson correlation coefficients (PCC), and Spearman correlation coefficients (SCC) between real and generated Protein produced by inClust+ and sciPENN were plotted.

## Discussion

In this paper, we describe a means to enhance inClust through adding the input-mask module and the output-mask module, and called the augmented version of model inClust+. We applied inClust+ to various datasets, ranging from multiple monomodal (unpaired) datasets, one or several multimodal (paired) datasets, and dataset containing multimodal data and monomodal data. In all cases, inClust+ demonstrated its capability to integrate, impute and generate data, and its performance was better than or comparable to the latest model. Firstly, inClust+ was used to integrate two monomodal data, namely, scRNA data and MERFISH data. It also could impute MERFISH data by extracting information from scRNA data. Then, inClust+ was used to integrate multimodal PBMC data containing scRNA data and scATAC data. The results show that it can’t only mix data between modalities, but also separate biological differences. Next, inClust+ was used to integrate multiple multimodal data (CITE) with batch effect. The results show that the difference between different modalities and the batch effect between different datasets can be simultaneously removed by inClust+. Finally, inClust+ was used to integrate data with both monomodal dataset and multimodal dataset. The results show that inClust+ could transfer label from multimodal data to monomodal data, and can complete the missing modality in monomodal data. In this study, we only use the paired data to test inClust+. The application scope of inClust+ is not limited to paired data, but also suitable for data with triple or more modalities.

Adding masks is a common way to enhance models in deep learning[27]. In inClust+, we augment our model through a pair of masks module (the input-mask module and the output-mask module). The flexible design and use of masks enable model to complete a series of tasks, which usually need to be completed by multiple models respectively. For example, inClust+ can utilize the common and dataset-specific genes for integration and imputation, as uniPort[18]. Masking makes things simple: the input-mask screens out common genes and the output-mask screens out common and dataset-specific genes of the corresponding data. Meanwhile, inClust+ could integrate multimodal dataset to achieve multi-domain translation, as cross modal autoencoder[21]. Input-mask and output-mask make inClust+ into two independent and related encoder-decoder combinations. Therefore, inClust+ can not only compress and reconstruct the data from the same modality, but also compress the data from one modality and reconstruct it into another modality, thus realizing cross modal translation. Furthermore, inClust+ could integrate multimodal datasets and monomodal dataset, transfer labels from multimodal data to monomodal data, and complete monomodal data into multimodal data by data generation, as sciPENN[25]. InClust+ refers to multimodal dataset to generate the data of missing modality in monomodal dataset. Generally speaking, as a model augmentation technology, adding a pair of masks to the model is not only limited to inClust, but also can be extended to deep learning models with similar encoder-decoder structures, such as scArches[28].

## Methods

### Data sets and preprocessing

#### Brain scRNA and MERFISH dataset

The mouse brains’ scRNA-seq and spatial transcriptomics datasets were obtained from Gene Expression Omnibus (GSE113576) [29] and Dryad repositories [https://datadryad.org/stash/dataset/doi:10.5061/dryad.8t8s248], respectively. Then, the data were preprocessed according to the method of Cao et al[18]. We obtained 30370 cells in scRNA and 64373 cells in MERFISH with 153 common genes.

#### Human PBMC paired multi-omics dataset

The single cell paired-omics dataset including DNA accessibility and gene expression comes from the publicly available dataset (PBMCs from C57BL/6 mice (v1, 150×150), Single Cell Immune Profiling Dataset by Cell Ranger 3.1.0. 10× Genomics, 2019), and were preprocessed according to the method of Cao et al[18]. We obtained 11259 cells with 2000 highly variable common genes across cells of all datasets.

#### Human PBMC and MALT CITE-seq dataset

The CITE-seq of human PBMC datasets was obtained from Gene Expression Omnibus (GSE164378)[30]. Then, we selected highly variable genes (HVGs) according to the method of Lakkis et al[25]. For gene, we used the scanpy 1.7.1 to normalize expression value[31]. Finally, for the PBMC dataset, we obtained 161748 cells with 1000 HVGs and 224 proteins. Cells from donor 7 (25827 cells) and donor 8 (26208 cells) were used in the integration experiment. In cross modal generation experiment, cells from donor 6 (20651 cells) and donor 7 (25827 cells) were used as multimodal dataset and scRNA data in donor 8 (26208 cells) was used as monomodal dataset.

### InClust+: overview

InClust+ is based on inClust, which integrates information from multiple sources and can work in supervised, semi-supervised and unsupervised modes[26]. The model is based on the variational autoencoder (VAE) and its variants.

### Architecture of inClust+

#### Input

InClust+ take 5 inputs, input 1 is the multimodal data and input 2 is the covariates information such as batch or modality. Input 3 is the label information, which is optional. Input 4 is the input-mask for screening out input. Input 5 is the output-mask for screening out output.

#### Input-mask module

The input-mask is a matrix as big as the input, with 0 or 1 in each element. The input is multiplied element-wise with the input-mask matrix to screen out the desired elements.

#### Encoder

The encoder is a three-layer neural network with decreasing dimensions (input_in_, h_1_, h_2_). Each layer uses a non-linear function as the activation function.

#### Latent sampling layer

Both parameters mean (μ_z_) and standard deviation (Σ_z_) are estimated from h_2_, using a neural network without activation function. The reparameterization trick was used for sampling latent variables Z_1_.

#### Embedding layer

The embedding layer embeds the auxiliary information (input2) into the latent space as a real-valued vector (E). For example, the unwanted covariates (e.g.: batch, donor) or auxiliary information that is represented by a one-hot vector or real-valued vector are embedded into a real-valued vector in latent space.

#### Vector arithmetic layer

The vector arithmetic is performed in the latent space. The estimated mean (μ_z_) would substrate (or add) the embedding vector E. The resulting vector Z_2_ retains the real biological information after removing the unwanted covariates or mixing the auxiliary information.

#### Classifier

The real-valued vector Z_2_ will pass through a neural network with softmax as the activation function. The output of the classifier is the output2.

#### Decoder

The decoder is a three-layer neural network with increasing dimensions (Z_1_, h_3_, output1). Each layer uses a non-linear function as the activation function.

#### Output-mask module

The output-mask is a matrix as big as the output, with 0 or 1 in each element. The output is multiplied element-wise with the output-mask matrix to screen out the desired elements.

### Pseudocode

1. screen out the input by masking: input_in_ = input_1_⊙input_4_(input-mask)
2. encode the input into latent space: h_2_ = relu(W_2_(relu(W_1_* input_in_)))
3. estimate the parameter: μ_z_ = W_μ_h_2_, Σ_z_ = W_Σ_h_2_ Reconstruction:
4. reparameterization trick: Z_1_ = μ_z_ + Σ_z_*ε ε ∼ N(0,1)
5. decode the latent space vector z into expression profile:

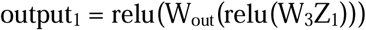
6. screen out the output by masking: output = output_1_⊙input_5_(output-mask) Clustering:

3: embed the auxiliary information: *E* = W_E_*input_2_

4: latent space vector arithmetic: Z_2_ = μ_z_ - *E* (or μ_z_ + *E*)

5: classification (clustering): output_2_ = softmax(W_C_Z_2_)

## Methods for Comparison

### uniPort

We applied the uniPort from the python package uniPort to data integration and imputation[18].

### sciPENN

We applied the sciPENN from the python package sciPENN to cross modal generation[25].

## Acknowledgments

This work was supported by grants from the National Key R&D Program of China [2021YFF1200904 to C.J.]; the Beijing Natural Science Foundation [Z210011 to C.J.]; the National Natural Science Foundation of China [32070795 to C.J. and 61673070 to J.Z.]

## Author contributions

L.W., J.C. and J.Z. envisioned the project. L.W. implemented the model. L.W. performed the analysis. L.W. wrote the paper. R.N., Y.C., A.W., H.Z., J.Z. and J.C. provided assistance in writing and analysis.

## Additional information

Correspondence and requests for materials should be addressed to J.C., J.Z. or L.W.

## Figure legend

**Figure S1.**
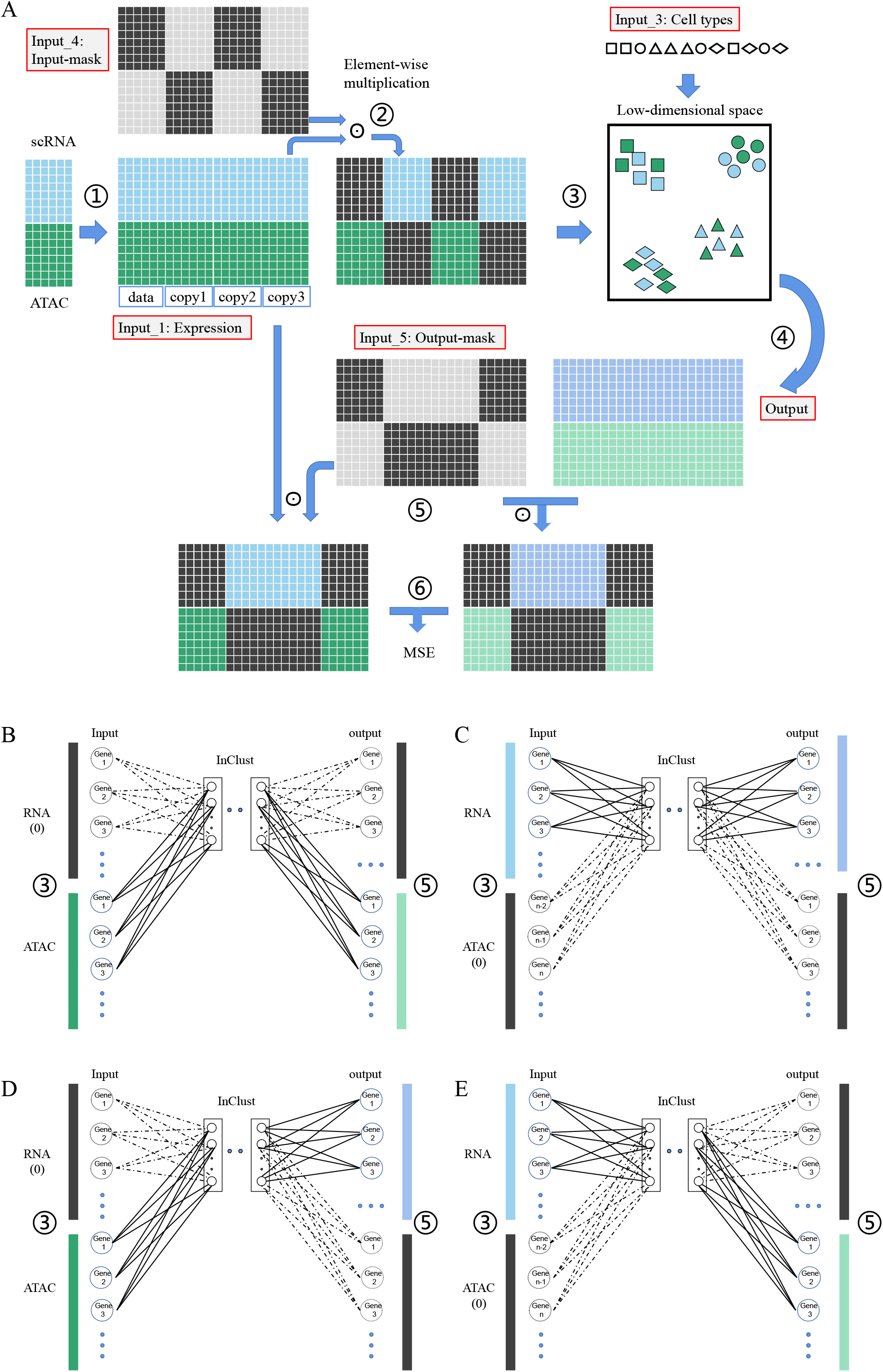
The diagram for integration of paired PBMC data (scRNA and ATAC) by inClust+ A. The workflow of inClust+ for integration of paired PBMC data (scRNA and ATAC). Training: ①Generation of the training dataset. To form a training dataset, the data of paired scRNA and ATAC data were duplicated 3 times, and concatenated together. ②Generation of the masked-input for the encoder in inClust. The training dataset multiplies element-wise with an input-mask matrix. The input-mask matrix is as big as the training dataset, and could be equally divided into four parts, each part being as big as the original paired dataset. In the first and the third part, the positions of scRNA data are filled with 0, and the positions of ATAC data are filled with 1. Alternatively, in the second and the fourth part, the positions of scRNA data are filled with 1, and the positions of ATAC data are filled with 0. The result of multiplication is the masked-input for the encoder, with alternate training data of scRNA and ATAC. ③Data encoding and data integration. The data from different modalities (scRNA or ATAC) in the masked-input are encoded by the different parts of encoder into the low-dimensional space, and integrated through the constraints of the cell type information. ④Reconstruction for both scRNA data and ATAC data. The decoder simultaneously outputs the reconstructed scRNA data and the reconstructed ATAC data. ⑤Generation of the mask-output for loss calculation. The output multiplies element-wise with an output-mask matrix. The output-mask matrix is as large as the output, and could be equally divided into four parts, each part being as big as the original paired dataset. In the first and the fourth part, the positions of scRNA data are filled with 0, and the positions of ATAC data are filled with 1. Alternatively, in the second and the third part, the positions of scRNA data are filled with 1, and the positions of ATAC data are filled with 0. The result of multiplication is the masked-output. ⑥Calculation of the loss for backpropagation. The MSE between masked-output and the masked-input is calculated as the loss. Data integration: After training, encoded low-dimensional representations from two modalities (scRNA and ATAC) are mixed together and clustered according to the cell types. B. The first training phase. In this stage, due to the influence of mask, only ATAC data is effective for input and output. Therefore, only the corresponding connections in the first layer (lower part) of the encoder and the last layer (lower part) of the decoder actually contribute to the training process. In short, inClust+ uses ATAC data to reconstruct ATAC data. C. The second training phase. In this stage, due to the influence of mask, only scRNA data is effective for input and output. Therefore, only the corresponding connections in the first layer (upper part) of the encoder and the last layer (upper part) of the decoder actually contribute to the training process. In short, inClust+ uses scRNA data to reconstruct scRNA data. D. The third training phase. In this stage, due to the influence of mask, only ATAC data is effective for input and scRNA data is effective for output. Therefore, only the corresponding connections in the first layer (lower part) of the encoder and the last layer (upper part) of the decoder actually contribute to the training process. In short, inClust+ uses ATAC data to reconstruct scRNA data. E. The fourth training phase. In this stage, due to the influence of mask, only scRNA data is effective for input and ATAC data is effective for output. Therefore, only the corresponding connections in the first layer (upper part) of the encoder and the last layer (lower part) of the decoder actually contribute to the training process. In short, inClust+ uses scRNA data to reconstruct ATAC data.

**Figure S2.**
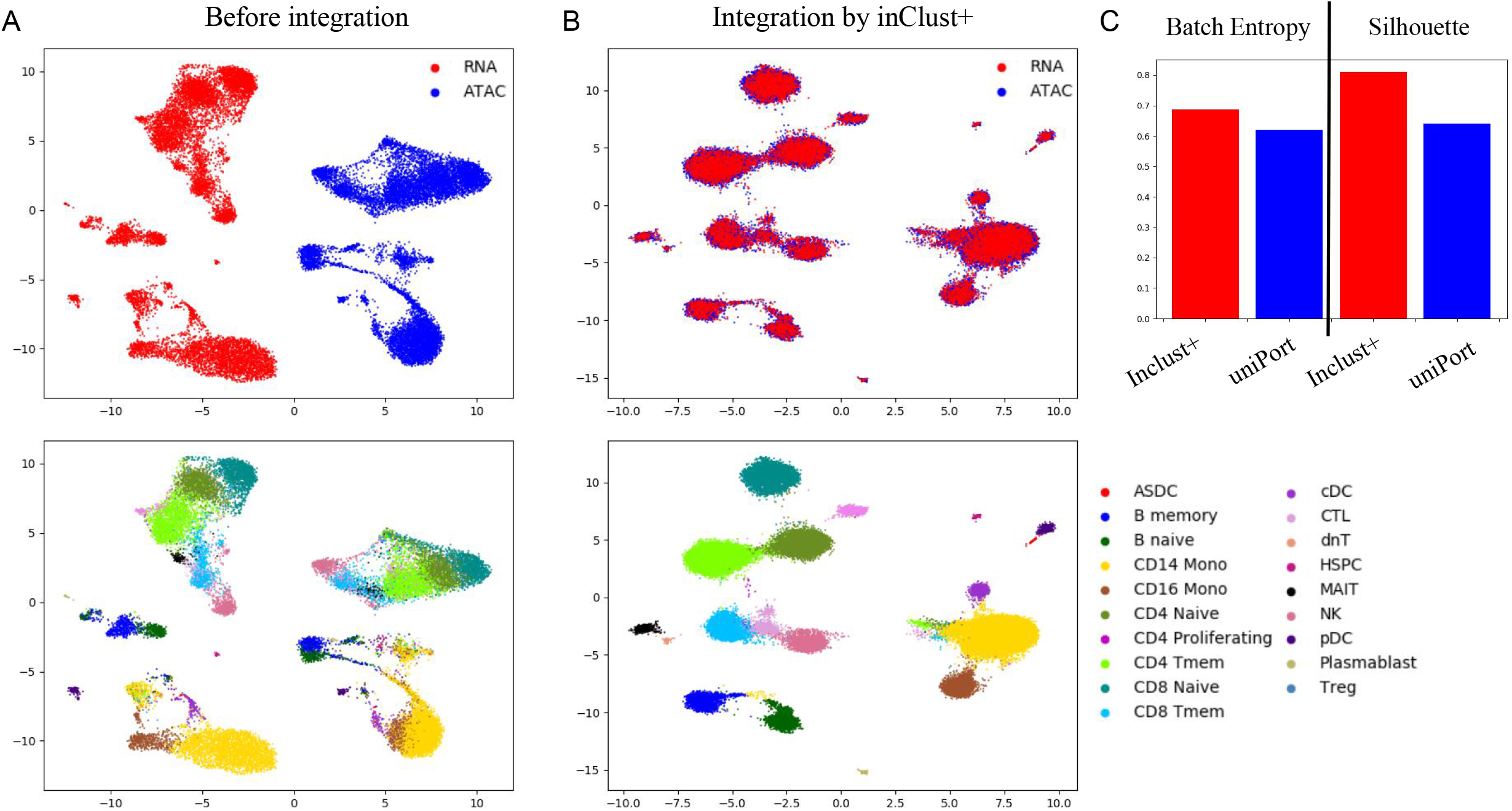
The results for integration of multiple multimodal (paired) data by inClust+. A. The UMAP plot of the scRNA and ATAC data colored by the covariate (top) and cell types (bottom). B. The UMAP plot of the low dimensional representations without covariate effects for the scRNA and ATAC data in inClust+ colored by the covariate (top) and cell types (bottom). C. The Batch Entropy score and the Silhouette coefficient measure from results of inClust+ and uniPort.

**Figure S3.**
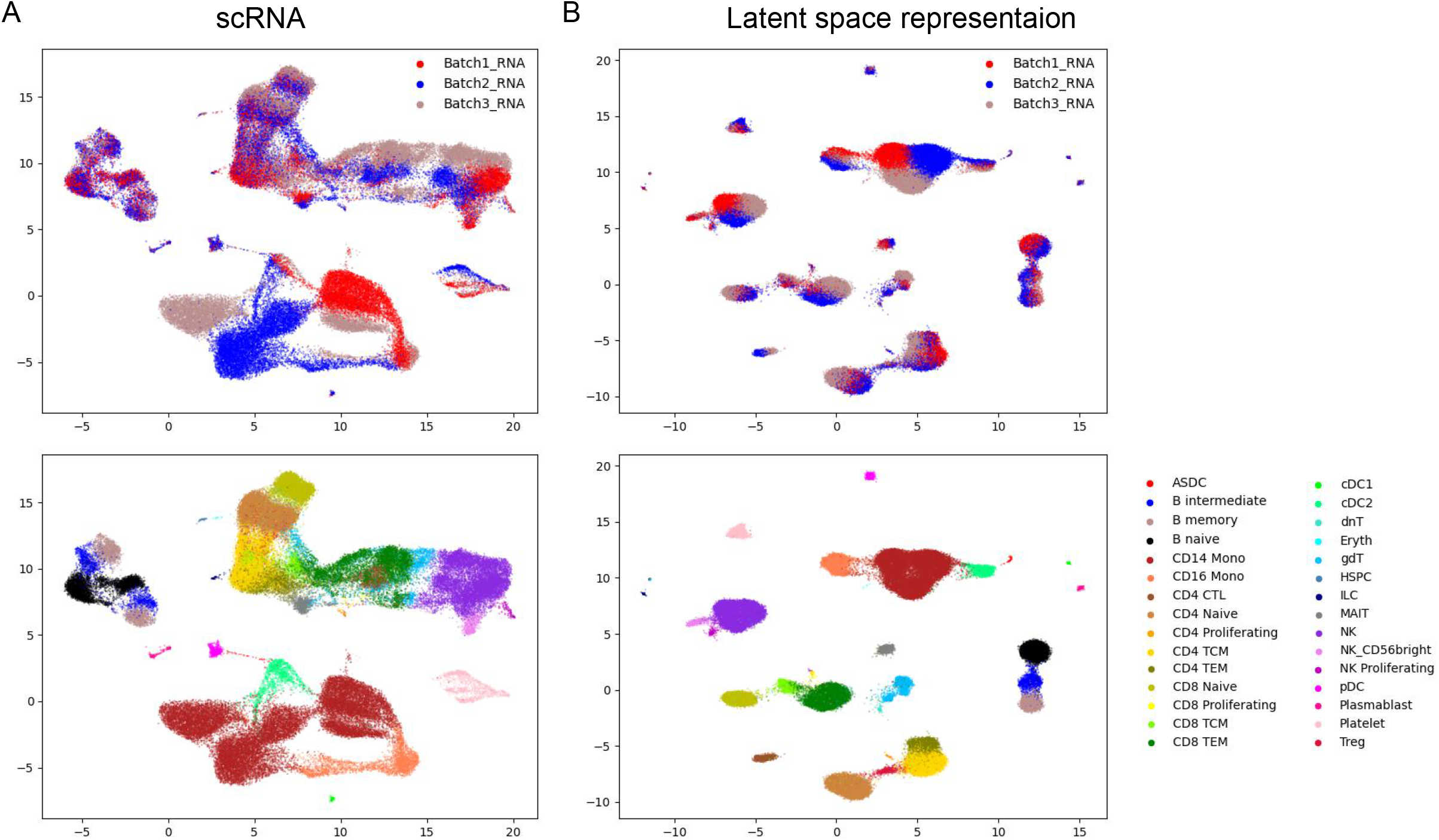
The results for integration of scRNA data in dataset with multimodal data and monomodal data A. The UMAP plot of the scRNA in the dataset colored by the batches (top) and cell types (bottom). B. The UMAP plot of the low dimensional representations with covariate effects for the scRNA data in inClust+ colored by the covariate (top) and cell types (bottom).

