## Supplementary figures and images for "InClust+: the multimodal version of inClust for multimodal data integration, imputation, and cross modal generation"

### Supplemental Figures

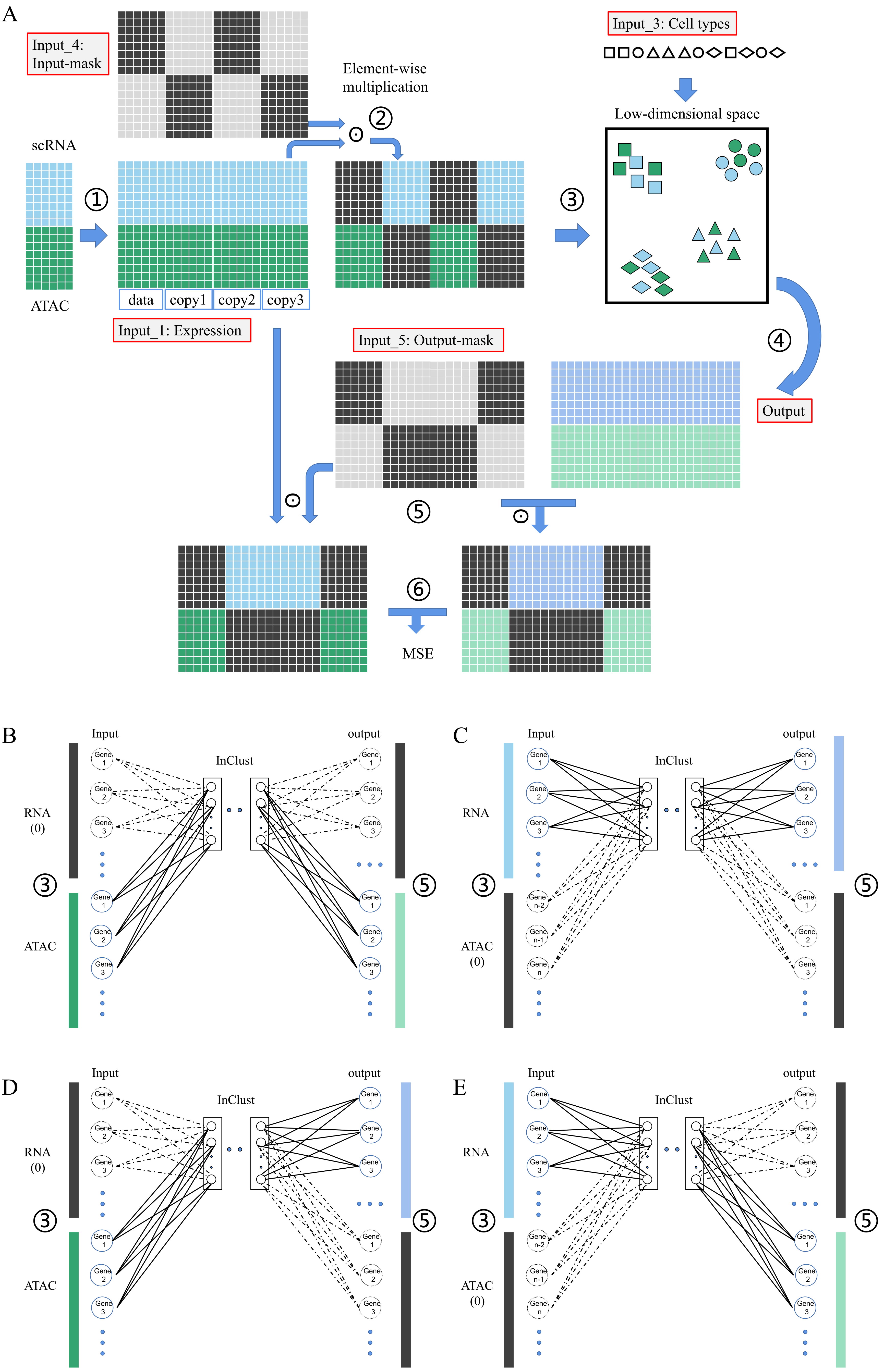

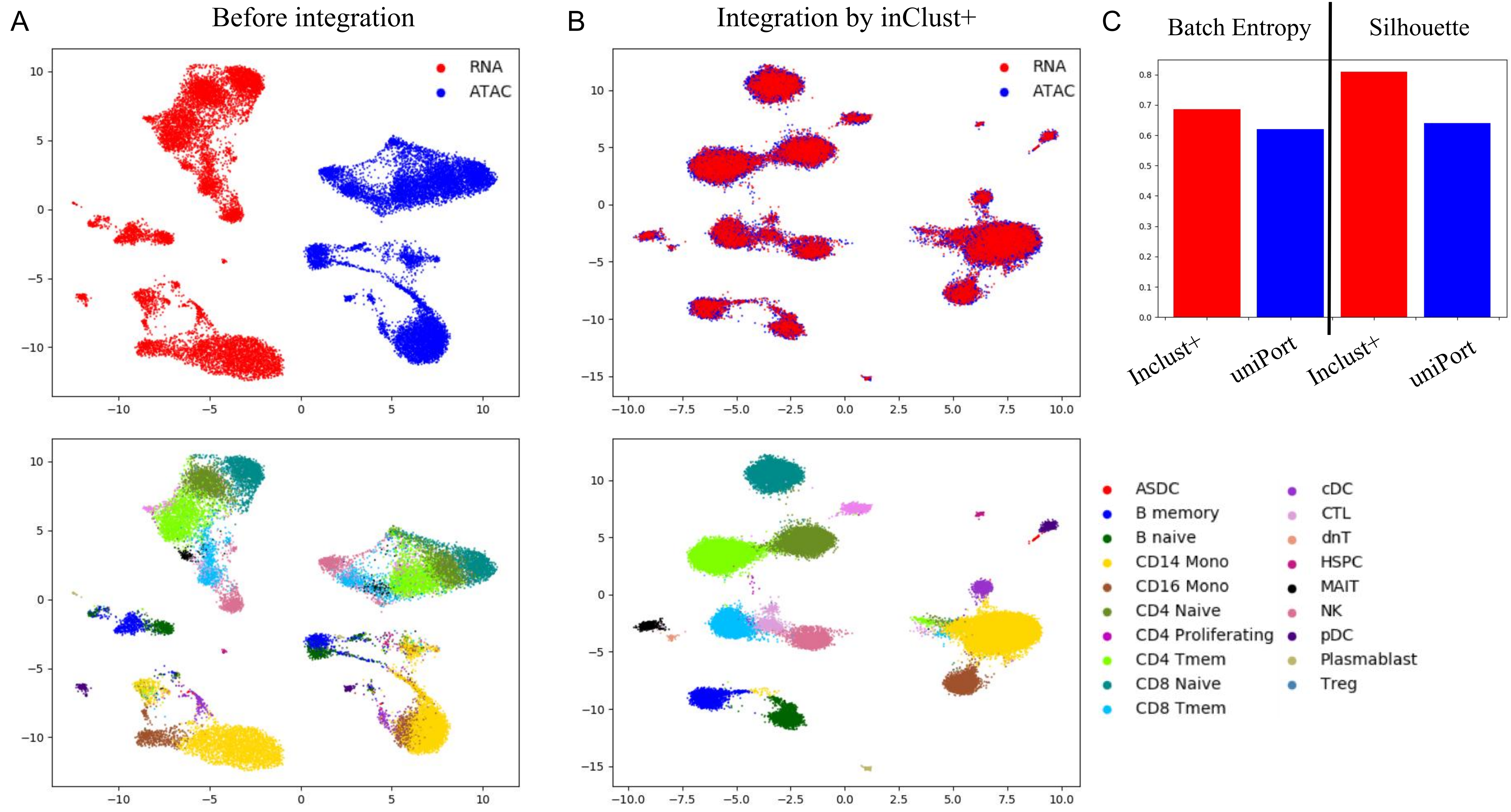

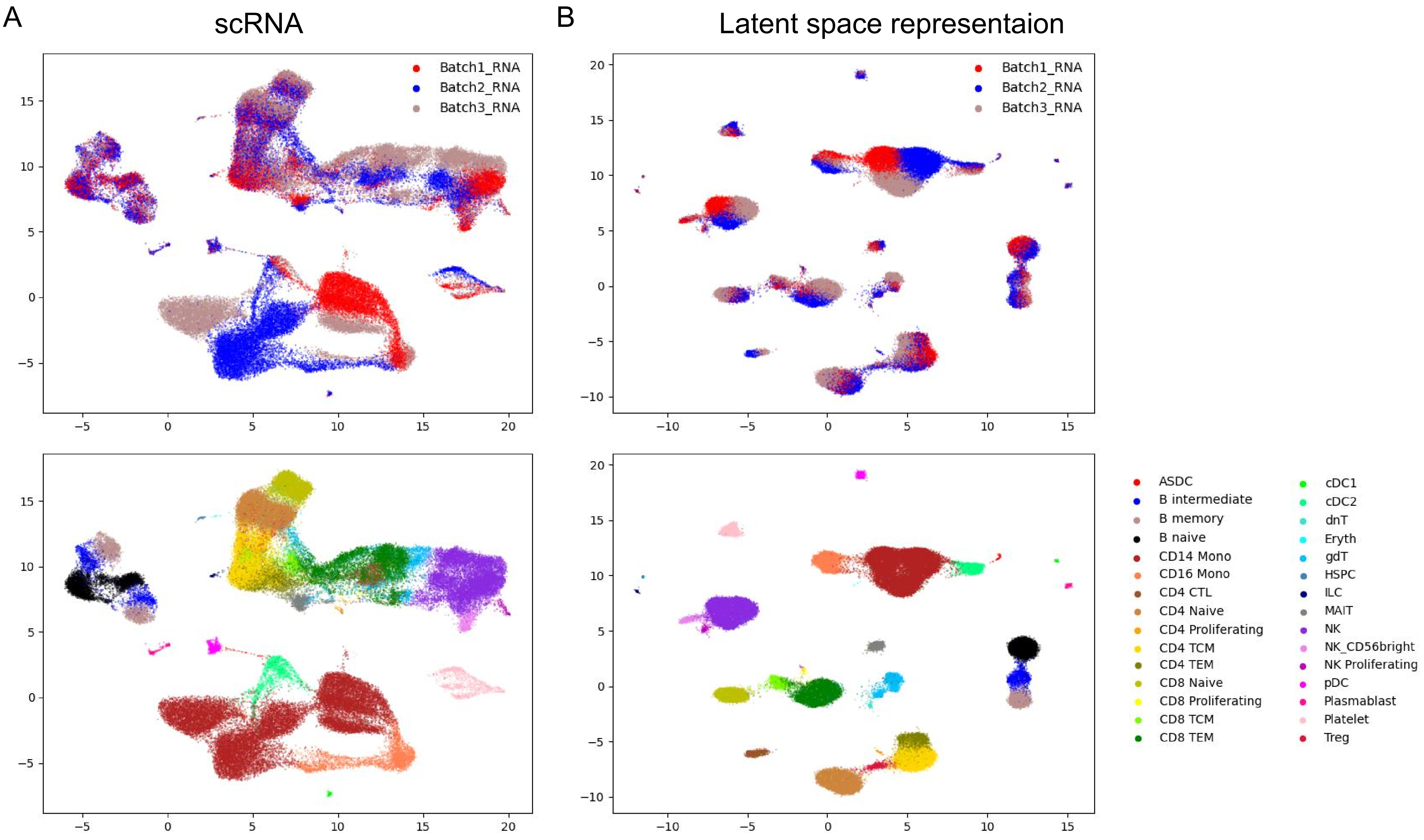
